# A reference bacterial genome dataset generated on the MinION™ portable single-molecule nanopore sequencer

**DOI:** 10.1101/009613

**Authors:** Joshua Quick, Aaron Quinlan, Nicholas J Loman

## Abstract

**Background:** The MinION™ is a new, portable single-molecule sequencer developed by Oxford Nanopore Technologies. It measures four inches in length and is powered from the USB 3.0 port of a laptop computer. By measuring the change in current produced when DNA strands translocate through and interact with a charged protein nanopore the device is able to deduce the underlying nucleotide sequence.

**Findings:** We present a read dataset from whole-genome shotgun sequencing of the model organism *Escherichia coli* K-12 substr. MG1655 generated on a MinION™ device during the early-access MinION Access Program (MAP). Sequencing runs of the MinION™ are presented, one generated using R7 chemistry (released in July 2014) and one using R7.3 (released in September 2014).

**Conclusions:** Base-called sequence data are provided to demonstrate the nature of data produced by the MinION™ platform and to encourage the development of customised methods for alignment, consensus and variant calling, de novo assembly and scaffolding. FAST5 files containing event data within the HDF5 container format are provided to assist with the development of improved base-calling methods. Datasets are provided through the GigaDB database http://gigadb.org/dataset/100102

## Introduction

Single molecule sequencing using biological nanopores was proposed nearly 20 years ago, but formidable technical challenges needed to be overcome before nucleotide sequence data could be reliably detected [1, 2, 3, 4, 5]. In Spring 2014, Oxford Nanopore Technologies released the first commercially-available nanopore sequencer to early-access customers. The MinIONT™ is no larger than a typical smartphone and can connect to and draw power from a laptop computer via its USB 3.0 interface. Sequence data is streamed as DNA fragments translocate through the pore, permitting real-time analysis on an Internet-connected laptop. Portable sequencing may open up new potential applications, for example near-patient testing and continuous environmental monitoring. We present the first bacterial genome data of the model organism *Escherichia coli* K-12 substr. MG1655 sequenced on the MinION™ during the MinION Access Program (MAP). Two flowcell chemistries, R7 (released July 2014) and R7.3 (released September 2014) were used. We anticipate this dataset will serve as a useful reference for the community to develop novel bioinformatics methods for this platform.

## Methods

### DNA extraction

*E. coli* K-12 MG1655 was streaked onto plate count agar and incubated for 48 hours at room temperature. Organisms were harvested using a L-shaped spreader and resuspended in 100 μl phosphate buffered saline (PBS). DNA extraction was performed using the Invisorb Spin Cell Mini Kit (Invitek, Birkenfeld, Germany) using the manufacturer’s protocol for extraction from serum or plasma.

### Sequencing library preparation

DNA was quantified using a Qubit fluorometer (Life Technologies, Paisley, UK) and diluted to 23.5 ng/μl. 85 μl was loaded into a G-tube (Covaris, Brighton, UK) and centrifuged at 5000 rpm for 1 minute before inverting the tube then centrifuging again for 1 minute. The fragmented DNA was end-repaired in a total volume of 100
μl using the NEBNext End-Repair module (NEB, Hitchin, UK). End-repair was performed as per manufacturer’s instructions except that the incubation time was reduced to 15 minutes. The resulting blunt-ended DNA was cleaned-up using 1.0x by volume AMPure XP beads (Beckman Coulter, High Wycombe, UK) according to the manufacturers instructions with the exception that 80% ethanol was used instead of 70%, and eluted in 25 μl molecular grade water. A-tailing was performed using the NEBNext dA-tailing module (NEB) in a total volume of 30 μl according to the manufacturers instructions with the exception that the incubation time was reduced to 15 minutes. We had concluded from previous experience that incubation times specified are unnecessarily long, and it is likely these incubation times could be further shortened.

### Sequencing library preparation: R7-specific

For the R7 chemistry run the Genomic DNA sequencing kit (*SQK-MAP-002*) (Oxford Nanopore Technologies, Oxford, UK) was used to generate a MinION sequencing library. To the 30 μl dA-tailed DNA, 50 μl Blunt/TA ligase master mix (NEB) was added in addition to 10 μl each of *adapter mix* and *HP adapter*. The reaction was left to proceed at room temperature for 10 minutes. The sample was cleaned-up using 0.4x by volume AMPure XP beads according to the manufacturers instructions with the exceptions that the kit supplied wash and elution buffers were used, and only a single wash was carried out. The sample was eluted in 25 μl of elution buffer. 10 μl of *tether* was added and incubated for 10 minutes at room temperature. Lastly, 15 μl of *HP motor* was added and incubated for 30 minutes at room temperature, giving a total volume of 50 μl library.

### Library preparation: R7.3-specific

In September 2014, an updated Genomic DNA sequencing kit (*SQK-MAP-003*) was released at the same time as a new set of flow cells termed R7.3. In this kit the *HP motor* is prebound to the hairpin adapter, eliminating the incubation step. To the 30 μl dA-tailed DNA, 50 μl Blunt/TA ligase master mix (NEB) was added in addition to 10 μl each of *Adapter Mix* and *HP adapter*, the reaction was left to proceed at room temperature for 10 minutes. The sample was cleaned-up using 0.4x by volume AMPure XP beads according to the manufacturer’s instructions with the exceptions that the kit-supplied wash and elution buffers were used, and only a single wash was used. The sample was eluted in 25 µl of elution buffer, leaving to incubate for 10 minutes before pelleting and removal of the library.

### Flowcell preparation

For each run new flowcell was removed from storage at 4°C and the protective packaging removed. The flowcell was fitted to the MinION™ device and held in place with supplied plastic screws to ensure a good thermal contact. 150 μl *EP buffer* was loaded into the sample loading port using a P1000 pipette and left for 10 minutes to prime the flowcell. The priming process was repeated a second time.

### Sample loading

Each library was quantified using the Qubit fluorometer. 100 ng (R7) or 350 ng (R7.3) of library was diluted into 146 μl using *EP buffer* and 4 μl *fuel mix* was added. The diluted library was loaded into the sample loading port of the flowcell using a P1000 pipette.

### Initiation of sequencing

A 72-hour (R7) or 48-hour (R7.3) sequencing protocol was initiated using the MinION control software, MinKNOW version 0.45.2.6 (R7) or 0.46.1.9 (R7.3). Read event data were base-called by the software Metrichor (version 0.16.37960) using workflow 1.0.3 (R7) or 1.2.2 rev 1.5 (R7.3).

### Data analysis

Read data was extracted from the native HDF5 format into FASTA using poretools[6]. Histograms of read length and collector's curves of reads were generated using the poretools *hist* and *yield plot* functions. Alignments were performed against the *E. coli* K-12 MG1655 reference sequence (accession U00096) using the LAST[7] aligner (version 475) with two sets of parameters, each of which were determined to give high mapping rates. Both LAST alignment settings use a gap opening penalty of 1 (option -a1) and a gap extension penalty of 1 (option -b1). Values of 1 (-q1) and 2 (-q2) were tried for the nucleotide substitution penalty parameters. Read percentage identity is defined as 100 * matches / (matches + deletions + insertions + mismatches). Fraction of read aligned is defined as (alignment length + insertions - deletions) / (alignment length + unaligned length deletions + insertions). Scripts used to generate alignments and plots are available at https://github.com/arq5x/nanopore-scripts

## Results

Datasets presented here have been deposited into the GigaDB resource at http://gigadb.org/dataset/100102

MinION reads can be classified into three types: template, complement and two-direction (2D). Template reads result from the first of two strands presented to the pore. Template strands are slowed down by a proprietary processive motor enzyme which is ligated to the leader adapter. Complement reads may be present if a hairpin has been successfully ligated. The shift from template to completement is recognised by an abasic site which produce a characteristic signal when in contact with the pore. The complement strand is slowed down a second enzyme termed the *HP motor*.

When both template and complement strands are sequenced these are combined by the base-calling algorithm to produce 2D reads, which may be divided into two types. *Normal 2D* are defined by having fewer events detected in the complement strand than in the template strand. This suggests that there was no *HP motor* bound to the hairpin to retard its process through the pore. *Full 2D* reads are defined as having more or equal complement events than template events. These are the optimal type of reads for analysis as they are of the highest quality (Figure 2).

**Figure 1.**
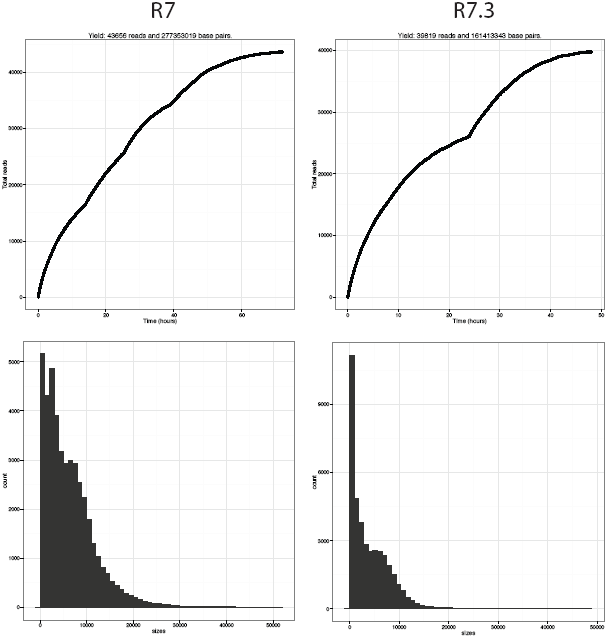
The top row plots demonstrate collector's curves of sequence reads over time measured in hours for the R7 and R7.3 run. The bottom row shows the histogram of read counts less than 50,000 bp in length for the R7 and R7.3 run.

**Figure 2.**
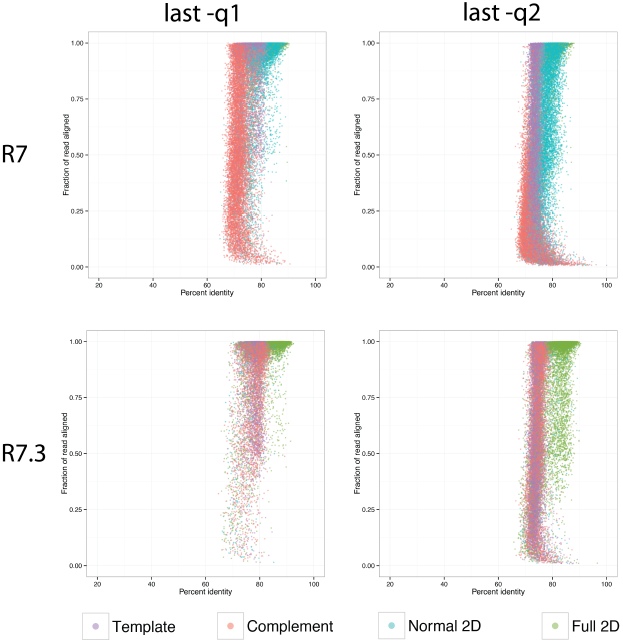
Alignment identity and completeness. Each plot reflects the alignment identity and the proportion of the read aligned for all 2D reads, as well as the underlying template and complement sequences. The top two panels reflect the alignment results for normal and full 2D reads from the R7 flowcell, and the bottom two panels reflect the R7.3 flowcell. Left panels employ a mismatch penalty of 1 and right panels reflect a mismatch penalty of 2. Overall, the lower mismatch penalty increases the identity and fraction of the read that aligned and this effect is greatest for full 2D reads.

The first run (R7) produced 43,656 template reads (272Mb), 23,338 complement reads (125 Mb) and 20,087 2D reads (131 Mb) of which 8% (10 Mb) were full 2D. The mean fragment length for 2D reads was 6,543. The second run (R7.3) produced 39,819 template reads (163 Mb), 18,889 complement reads (84 Mb), 11,823 2D reads (64.53 Mb) of which 86% (55.68 Mb) were full 2D (Table 1 and Table 2). The mean fragment length for 2D reads was 5,458.

**Table 1.** Yields for each nanopore run in reads.

|  | File | files | Template | Complement | All 2D | Full 2D |
| --- | --- | --- | --- | --- | --- | --- |
| 1 | R7 Non-overnight | 47195 | 43656 | 23338 | 20087 | 1598 |
| 4 | R7.3 | 42316 | 39819 | 18889 | 11823 | 9563 |

**Table 2.** Yields for each nanopore run in megabases.

|  | File | Template | Complement | All 2D | Full 2D |
| --- | --- | --- | --- | --- | --- |
| 1 | R7 Non-overnight | 272.07 | 125.03 | 131.44 | 9.56 |
| 4 | R7.3 | 162.68 | 84.35 | 64.53 | 55.68 |

2D reads contain information from both template and complement strands. Therefore a non-redundant dataset would consist of 2D reads plus any remaining template or complement reads which did not get turned into 2D reads.

We compared the alignment characteristics of full 2D sequences, as well their underlying template and complementary strand sequences, using the LAST aligner with two different parameter settings (see Methods). We observe a marked increase in the identity and the proportion of the read aligned when employing equivalent mismatch, gap introduction, and gap extension penalties (Figure 2). The effect is particularly pronounced for the more accurate full 2D reads, and this effect is consistent for both R7 and R7.3. Notably, the proportion of Full 2D reads was substantially (22.6% vs. 3.9%) higher in the R7.3 run, suggesting the possibility that future improvements to the chemistry will increase the yield of the highest quality and most biologically informative reads.

## Discussion

We show that the MinION™ is able to sequence entire bacterial genomes in a single run. Further work is required to determine appropriate algorithms for common secondary analysis tasks such as variant calling and de novo assembly. We anticipate and hope this dataset will help stimulate the development of novel methods for handling Oxford Nanopore data.

## Acknowledgements

### Competing interests

NJL and AQ are part of the Oxford Nanopore MinION Access Programme. (MAP). Oxford Nanopore have contributed free of charge early-access reagents in support of the data presented in this manuscript.

### Author’s contributions

JQ performed the DNA extraction, library preparation and sequencing and assisted in the data analysis. AQ performed the read alignment analysis. NJL performed the data analysis. NJL and JQ drafted the manuscript. All authors approved the final manuscript.

### Acknowledgements

NJL is funded by a Medical Research Council Special Training Fellowship in Biomedical Informatics. JQ is funded by the National Institute for Health research (NIHR) Surgical Reconstruction and Microbiology Research Centre (partnership between University Hospitals Birmingham NHS Foundation Trust, the University of Birmingham and the Royal Centre for Defence Medicine). The views expressed are those of the author(s) and not necessarily those of the NHS, the NIHR or the Department of Health. ARQ was supported by the NIH AQ7 (NGHRI; 1R01HG006693-01). Funders had no role in the design or carrying out of this work. The Medical Research Council Cloud Infrastructure for Microbial Genomics (CLIMB) platform was used for data analysis.

